# Learning a tactile sequence induces selectivity to action decisions and outcomes in the mouse somatosensory cortex

**DOI:** 10.1101/2020.04.17.037143

**Authors:** Michael R. Bale, Malamati Bitzidou, Elena Giusto, Paul Kinghorn, Miguel Maravall

## Abstract

Sequential temporal ordering and patterning are key features of natural signals used by the brain to decode stimuli and perceive them as sensory objects. To explore how cortical neuronal activity underpins sequence recognition, we developed a task in which mice distinguished between tactile ‘words’ constructed from distinct vibrations delivered to the whiskers, assembled in different orders. Animals licked to report the presence of the target sequence. Mice could respond to the earliest possible cues allowing discrimination, effectively solving the task as a ‘detection of change’ problem, but enhanced their performance when deliberating for longer. Optogenetic inactivation showed that both primary somatosensory ‘barrel’ cortex (S1bf) and secondary somatosensory cortex were necessary for sequence recognition. Two-photon imaging of calcium activity in S1bf layer 2/3 revealed that, in well-trained animals, neurons had heterogeneous selectivity to multiple task variables including not just sensory input but also the animal’s action decision and the trial outcome (presence or absence of a predicted reward). A large proportion of neurons were activated preceding goal-directed licking, thus reflecting the animal’s learnt response to the target sequence rather than the sequence itself; these neurons were found in S1bf as soon as mice learned to associate the rewarded sequence with licking. In contrast, learning evoked smaller changes in sensory responses: neurons responding to stimulus features were already found in naïve mice, and training did not generate neurons with enhanced temporal integration or categorical responses. Therefore, in S1bf sequence learning results in neurons whose activity reflects the learnt association between the target sequence and licking, rather than a refined representation of sensory features.

## Introduction

Natural sensory signals unfold over time, and their temporal patterning is inherent to their identity. Being sensitive to this patterning allows sensory systems to identify known stimuli, detect new or unexpected stimuli, and distinguish between objects. Thanks to this capacity we can simultaneously recognize a favourite song playing on the radio and the identity of a family member from the cadence of their steps walking towards us. How is this ability underpinned by neuronal responses?

Within sensory modalities such as touch, the spiking responses of early sensory neurons faithfully relay temporally patterned signals to the brain for later integration and decoding [1-10]. Decoding such patterns could be facilitated by the known biophysical properties of central neurons and synapses. Neurons can become sensitive to specific spatiotemporal patterns of synaptic input [11, 12]; in vitro networks of neurons can intrinsically encode temporal input sequences [13, 14]; and synapses mediating thalamocortical input can have diverse temporal filtering properties [15]. However, how these capacities relate to sensory sequence learning in a living animal is unknown. How does neuronal activity in vivo distinguish between relevant sequences? Do neurons become categorically sensitive to a specific sequence as the result of learning?

We recently developed a tactile GO/NOGO discrimination task to test mouse and human capacities for sequence recognition [16]. Mice learned to recognize patterns of noisy, initially meaningless whisker vibrations whose amplitude was modulated in a specific sequence (an amplitude modulation envelope) lasting several hundred milliseconds. The target sequence differed from others only in the ordering of segments of different amplitude. Mice reached a level of discrimination performance comparable to humans, and seemed to base this performance on detecting cues (changes or transitions in the vibration pattern) that were differentially present in the target sequence as compared to others [16].

In the rodent tactile whisker system, sensory information first reaches the cortex through the ‘barrel field’ of the primary somatosensory cortex (S1bf), the main cortical target for direct somatosensory input from the thalamus [17]. Sensory-guided tasks mediated by whisker input can be carried out without cortical participation [18, 19], and recent results have highlighted that the flow of sensory information through cortical pathways is task-dependent [20]. Moreover, although neurons in S1bf encode well-defined stimulus features, in naïve animals they do not integrate sensory information over time [21-26]. Here, we determined whether S1bf and successive cortical processing stages are needed to solve the elementary sequence recognition task, and whether neuronal responses in S1bf become selective to the target sequence as a result of learning.

## Results

### Recognition of elementary tactile sequences in mice

We set out to determine the changes in cortical neuronal responses that result from learning to recognize a sequential tactile pattern. To this end, we trained head-fixed mice to respond selectively to a target sequence of vibrations delivered to the whiskers (Figure 1). In this GO/NOGO discrimination design, target and non-target sequences differed in the order of their (initially meaningless) elements. Stimulation was delivered to multiple whiskers (Methods).

**Figure 1.**
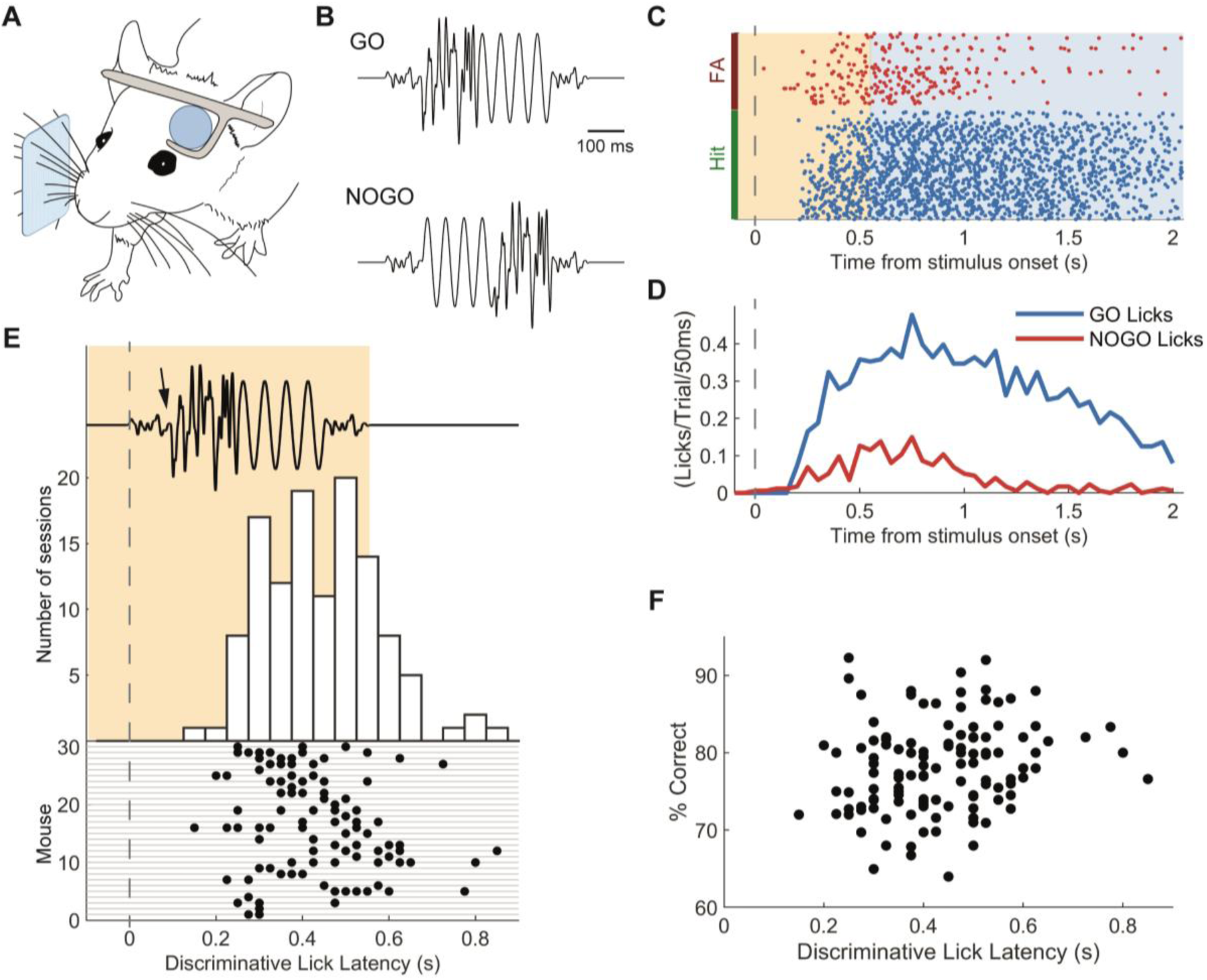
Recognition of elementary tactile sequences in mice. (**A)** Diagram of experimental set-up. Sensory stimulation was delivered to head-fixed mice via the whiskers. **(B)** The GO and NOGO stimulus sequences consisted of four segments, and differed in that the order of the central segments was switched. **(C)** Raster plot of licks on GO (hit) and NOGO (false alarm) trials for an example session. Shading shows stimulus presentation period. **(D)** Histogram of licks on GO and NOGO trials for the same example session. The time at which traces diverge is termed the discriminative lick latency. **(E)** Histogram and raster plot showing discriminative lick latency over the course of a session. Raster distinguishes data for all mice (each mouse, one row) and sessions (each session, one data point). Time is relative to start of stimulus sequence (top). Arrow shows time when target GO sequence (shown here) diverges from NOGO sequence and can first be distinguished. **(F)** Performance (% correct) plotted against discriminative lick latency across all sessions.

Building on our earlier finding that mice learn to recognize tactile sequences constructed as a concatenation of noise segments [16], we used a simple sequence design in which each stimulus consisted of a tactile ‘word’ (Figure 1B). Each segment within the ‘word’ comprised either filtered noise (with different amplitudes) or sinusoidal stimulation. A trial began with a ‘stimulus presentation period’ lasting 550 ms, in which the sequence was delivered. Mice were not rewarded or punished for licking during this period. At the end of this stimulation period followed a ‘response period’ (1.5 s) where mice needed either to lick or refrain from licking, depending on stimulus sequence. Following a GO sequence, if mice licked during the response period (hit trial) they received a water reward; if they failed to lick (miss trial) the next trial began as normal. Following a NOGO sequence, if mice correctly withheld licking during the response period (correct rejection trial) the next trial began as normal; if they licked (false alarm trial) the next trial was delayed by 2-5 s, with the duration set depending on mouse thirst and impulsiveness.

After training, head-fixed mice readily learned to associate one specific sequence with licking for a water reward (Figure 1C, D). Animals took 2-26 training sessions to reach 70% correct discrimination (mean 6.1 sessions, SD 5 sessions) and achieved a mean of 80% performance in their best-performing session (SD 7%, n = 42 animals).

What cues did mice use to recognize the learnt sequence? An ideal observer would be able to distinguish the identity of the target sequence immediately upon the first transition between its constituent segments, as this was the moment at which GO (target) and NOGO (non-target) sequences diverged from each other (Figure 1B). In our design, mice were allowed to lick during the sequence presentation period without incurring reward or punishment (Methods), and this allowed us to measure their freely varying response (lick) times. For each session, we determined the time at which lick rates on GO (hit) trials diverged significantly from those on NOGO (false alarm) trials. This gave an upper-bound estimate for when the mouse, on average, reached its decision as to sequence identity in that session (Methods). We term this measure ‘discriminative lick latency’ (DLL) (Methods; Figure 1D).

Across a data set of 30 mice and 120 sessions, DLL varied by animal and session (Figure 1E). The DLL was sometimes short enough to suggest that the animal made its decision immediately upon detecting the earliest possible GO cue; but in other sessions, the value of the DLL suggested longer deliberation (range 150-850 ms, mean 432 ms). Both strategies – instantaneous response or deliberation – could potentially lead to high performance, depending on conditions. An ideal instantaneous detector under noiseless conditions would be able to identify the target immediately upon the first transition (Figure 1E, arrow): in this scenario slower responses would imply no gain in performance, and might even be a signature of impaired performance in a poor learner. This situation would be reflected either in an absence of correlation between DLL and performance, or in longer DLLs corresponding to lower performance. On the other hand, under conditions where whisker stimulation is noisy and the identity of the sequence might become clearer over time, it could be beneficial for mice to be able to integrate sensory information for longer in order to do better. In this situation, longer DLLs would correspond to higher performance. We found that the latter applied: sessions with greater integration or deliberation, as measured by a longer DLL, correlated with higher performance (Figure 1F; 120 sessions; r = 0.20, p = 0.028, Pearson correlation; t = 2.41, p = 0.017, mixed-effects model with mouse ID as random factor).

These results demonstrate that mice readily learned to associate a specific tactile sequence with a water reward. Animals were often able to discriminate quickly, consistent with an ability to focus on detecting the earliest cues that allowed discrimination of the target sequence. However, performance on the task tended to be higher when animals took longer, consistent with deliberating or accumulating sensory evidence over tens to hundreds of milliseconds.

### Somatosensory cortex carries sensory information needed for sequence recognition

To track the flow of neuronal activity through early cortical stages during task performance and determine which stages were needed for sequence discrimination, we trained mice expressing channelrhodopsin in cortical GABAergic interneurons (VGAT-ChR2-EYFP; [27]). Once mice had achieved 75% correct performance during a session, we began running sessions combining optogenetics with behaviour. We suppressed activity in specific, stereotaxically defined regions of dorsal cortex throughout the stimulus presentation period (from 50 ms before stimulus onset to 50 ms after offset), illuminating the cortical surface with a blue laser (Methods). Laser-ON and laser-OFF trials were interspersed, with laser-ON comprising a randomly chosen subset (20%) of all trials. In addition to S1bf (coordinates: 1 mm anteroposterior from bregma [AP], 3 mm mediolateral [ML]), we selected the following regions for optogenetic suppression, all of which receive direct projections from S1bf: secondary somatosensory cortex (S2; 1.2 mm AP, 4.2 mm ML); posterior parietal cortex (PPC; 2 mm AP, 1.7 mm ML); and whisker primary motor cortex (wM1; 1.1 mm AP, 0.9 mm ML). S1bf and S2 are part of the ascending cortical pathway for tactile input. PPC is a centre for multisensory sensorimotor integration and its activity has been shown to accumulate sensory evidence over time and reflect history biases [28-30]. M1 is a centre regulating the learning and deployment of newly relevant motor responses, but can also accumulate sensory evidence over time [31] and acts as a goal-directed modulator or inhibitor, rather than just a generator, of motor actions [32].

We found that optogenetic suppression centred over either S1bf or S2 significantly decreased lick response rates (the percentage of trials with a lick response; Figure 2B, C) (S1bf: 4 mice, 10 sessions, F(1,36) = 97.8, p < 10^−11^; S2: 3 mice, 11 sessions, F(1,40) = 308, p < 10^−19^, both ANOVA 2-way test). This decrease in response rate affected both GO and NOGO trials (Figure 2B, C), although it was greater on GO trials (S1bf: F(1,36) = 7.40, p = 0.01; S2: F(1,40) = 45.7, p < 10^−7^). That the decrease in response rate with laser ON occurred both on GO and NOGO trials indicated that this form of optogenetic manipulation was not disturbing a specific representation of the GO target sequence; rather, it likely interfered with the overall flow of sensory information through S1bf and S2.

**Figure 2.**
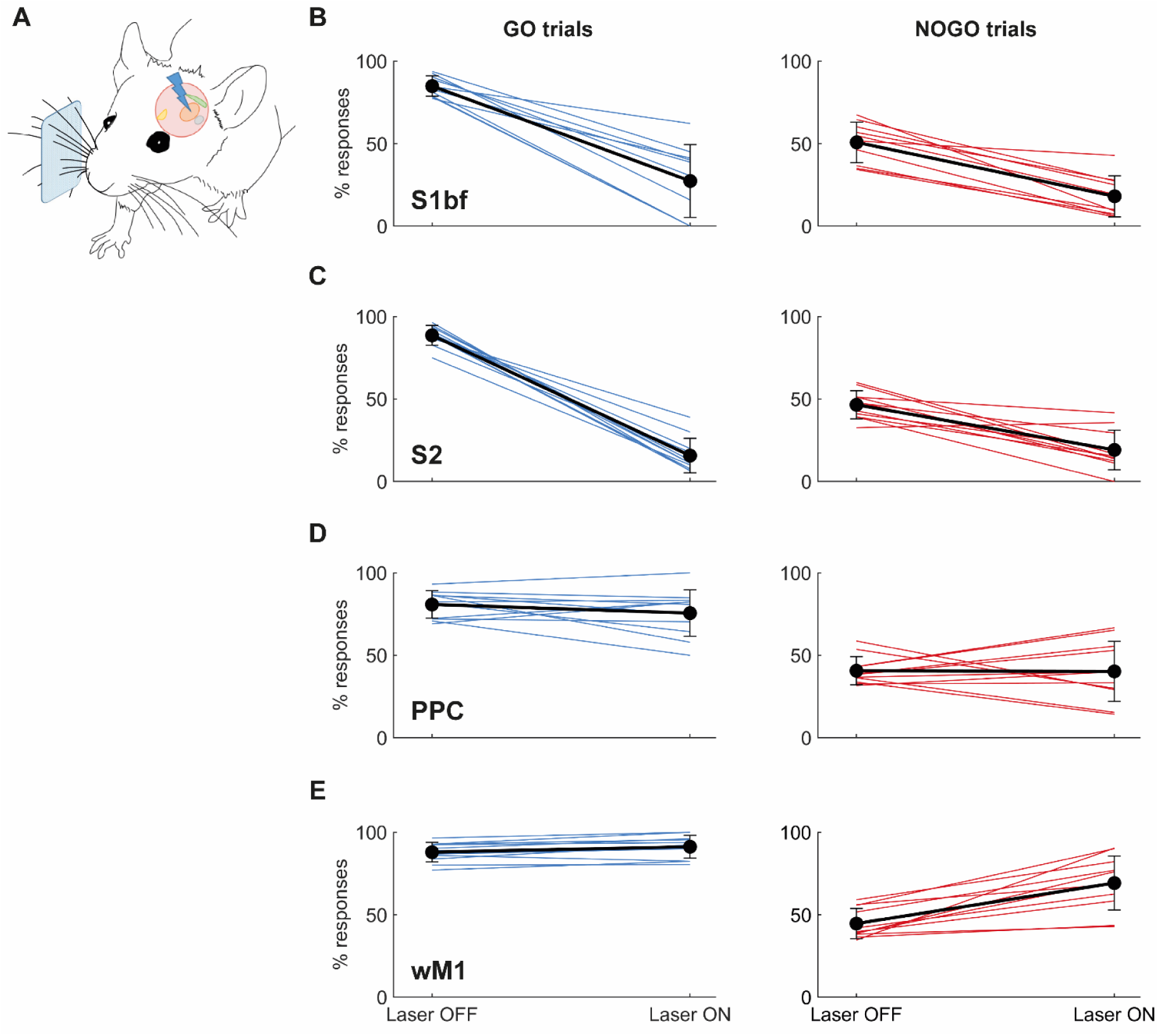
Tracking the participation of different cortical regions with optogenetic activity suppression. **(A)** Diagram of experimental set-up. Cortical areas were illuminated with a blue laser during trial performance. **(B)** Effects of S1bf suppression on performance (lick response rates) on GO (left) and NOGO trials (right). **(C)** Effects of S2 suppression. **(D)** Effects of PPC suppression. **(E)** Effects of wM1 suppression. Black lines: means across sessions; error bars: SD; coloured lines: individual sessions.

We also considered the alternative possibility that S1bf or S2 inactivation did not suppress sensory information needed for the decision but caused a nonspecific decrease in the probability or speed of elicited motor actions, leading to the reduced response rate. If manifested as an overall scaling down of licking probability, such an effect would still result in a lower rate of false alarms than hits. However, two lines of evidence speak against this. First, on laser-ON trials there was no significant difference between the probability of responding to GO and NOGO stimuli, so that discrimination fell to random levels (Figure 2B, C; d’ upon laser stimulation did not differ significantly from 0: S1bf inactivation, p = 0.967; S2 inactivation, p = 0.880, both Wilcoxon signed rank test). Second, there was no evidence for lick responses becoming slower: median latency to first lick was not significantly different on laser-ON and laser-OFF trials (S1bf inactivation, p = 0.578; S2 inactivation, p = 0.365, both Wilcoxon signed rank test).

These results indicate that the sensory signals necessary to recognise a sequence of vibrations delivered to the whiskers were routed through S1bf and S2, with optogenetic suppression causing effects consistent with a loss of sensory input to decision-making stages. Inhibition of either area had a similar effect, consistent with a serial flow of sensory information or with an S1bf-S2 loop activated in series [33-35]. As a control for the specific contribution of S1bf and S2 activity to tactile sequence recognition, we interfered optogenetically with these areas in animals that had learned to base sequence recognition on acoustic rather than tactile cues: in this case, inhibition did not affect lick response behaviour (2 mice, 2 sessions per mouse; data not shown).

Suppressing PPC had no detectable effect on response rate (Figure 2D) (3 mice, 11 sessions, F(1,40) = 0.52, p = 0.475, ANOVA 2-way test). Suppressing wM1 disinhibited lick responses particularly on NOGO trials, i.e. those in which mice had been trained selectively to avoid licking (Figure 2E) (3 mice, 11 sessions, F(1,40) = 19.7, p < 10^−4^, ANOVA 2-way test; interaction between trial type and effect of laser, F(1,40) = 11.5, p = 0.0016). These findings show that the sensory information necessary for learnt sequence recognition was carried by S1bf and S2 but not PPC or wM1.

### Neuronal responses in well-trained S1bf during sequence discrimination reflect heterogeneous task variables and learnt associations

To uncover the nature of neuronal responses in primary sensory cortex during sequence recognition, we used two-photon imaging in layer 2/3 of mice performing the task (Figure 3). Mice were Thy1-GCaMP6f animals expressing calcium indicator in cortical excitatory neurons [36] (Methods; n = 7 mice). For each neuron in a session, we extracted the differential fluorescence (ΔF/F_0_) time series on every trial and computed the average ΔF/F_0_ response profile parsed by trial outcome (hit, miss, false alarm, correct rejection), with reference to the beginning of the trial. We also computed the average ΔF/F_0_ response locked to the time of the first lick on hit and false alarm trials. This visualization allowed us to explore the relationship between trial type and changes in fluorescence (Figure 3C).

**Figure 3.**
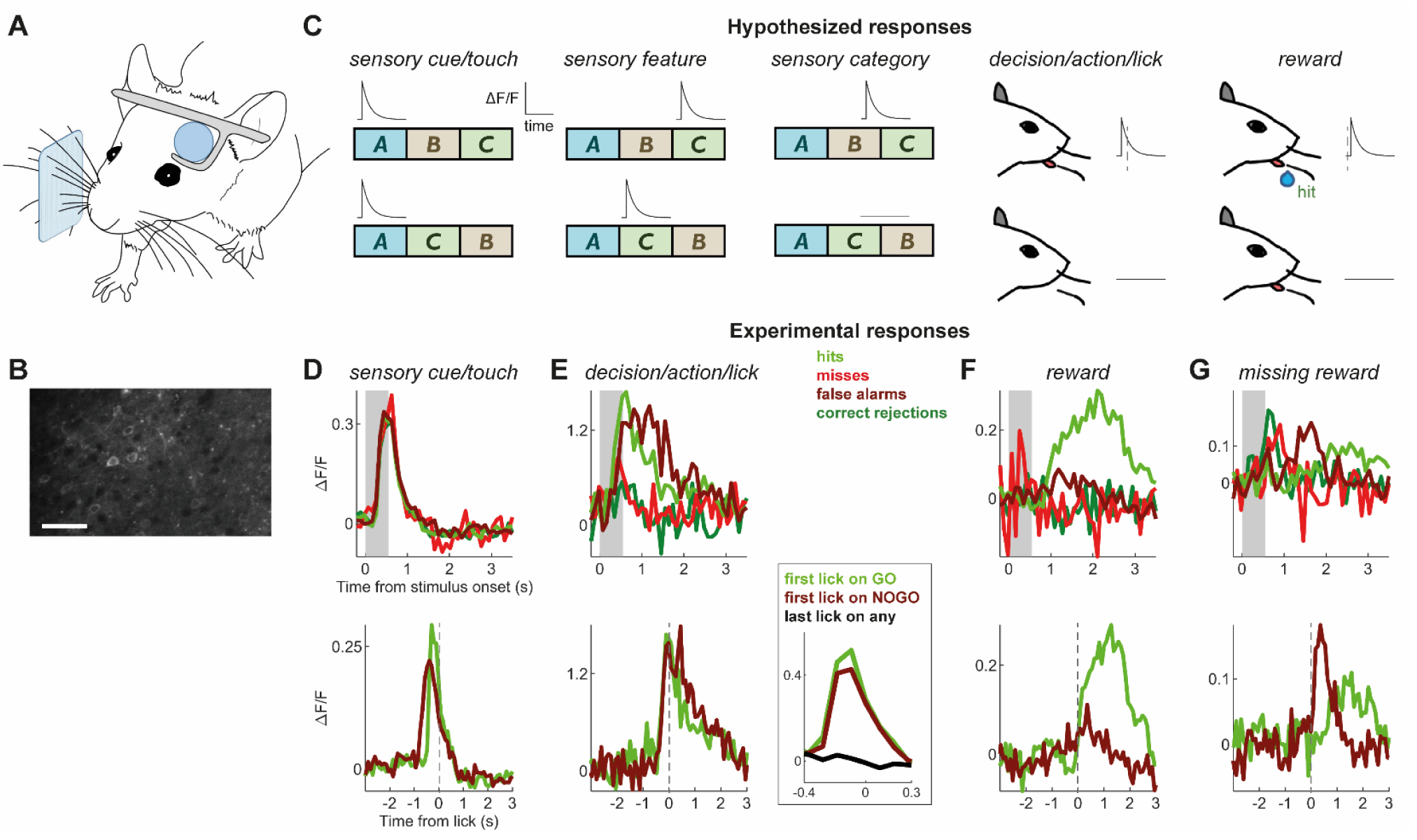
Neuronal responses in well-trained S1bf reflect heterogeneous task variables and learnt associations. **(A)** Diagram of experimental set-up. Two-photon imaging was carried out while animals performed the task. **(B)** Example of field of view (scale bar: 100 μm). **(C)** Hypothetical expected responses. Diagrams showing the ΔF/F_0_ responses presented by hypothetical neurons reflecting one of the following: sensory cue presentation; a specific sensory feature (stimulus segment C in the example); sensory category (ABC vs ACB); the prediction of the decision to lick; or the presence of a reward. In the last two cases, note the different timing of the response relative to the lick (dashed line). **(D-G)** Actual experimental responses. **(D)** Example experimental neuron responsive to sensory cue. Top, mean ΔF/F_0_ relative to stimulation time (shaded region: stimulus presentation period). Bottom, mean ΔF/F_0_ relative to first lick (dashed line). **(E)** Example neurons predictive of licking. Left panels, one neuron; right panel, different neuron highlighting the difference between responses to first lick on GO or NOGO trials and last lick on any trial. **(F)** Example neuron responsive to reward on hit trials. **(G)** Example neuron responsive to reward prediction error on false alarm trials.

Based on the notion that S1bf is an area that primarily provides sensory information to higher stages in a behavioural hierarchy for sensory-guided decision making, we expected our data to be dominated by neurons representing features of sensory stimulation sequences, potentially including neurons categorically selective to the overall identity of the learnt sequence (termed ‘sensory category’ neurons and responding only on GO trials, Figure 3C).

In the event, neurons in S1bf of well-trained mice showed remarkably heterogeneous selectivity to multiple task variables including the onset of sensory input (‘sensory cue’ neurons, Figure 3C, D), the animal’s decision to act with a lick (‘decision/action/lick’, Figure 3C, E) and the subsequent outcome of the trial including the presence or absence of expected rewards (‘reward’, Figure 3C, F, G). The most frequently found types (Table 1) were ‘sensory cue’ neurons responding to the onset of stimulation (Figure 3D) and ‘decision/action/lick’ cells activated just before goal-directed licking, the animal’s learnt response to the target sequence (Figure 3E). We note that GO and NOGO sequences shared a common initial segment, so that a neuron sensitive to stimulus onset and with a strongly adapting response to sustained stimulation would ‘view’ the sequences as being identical; many layer 2/3 neurons labelled as ‘sensory cue’ in the present task would likely appear as touch sensitive neurons in situations where an animal encounters objects with its whiskers [37-39].

**Table 1.** Proportions of visually identified neurons responding to sensory and task variables in well-trained S1bf.

| Sensory cue | Sensory feature selective | Sensory category | Decision/action predictive (during stimulus period) | Decision/action predictive (during response period) | Licking | Reward delivery and error | Unclassified |
| --- | --- | --- | --- | --- | --- | --- | --- |
| 53<br>(17%) | 0 | 1<br>(0.32%) | 70<br>(22%) | 5<br>(1.6%) | 43<br>(14%) | 7<br>(2.2%) | 136<br>(43%) |

Action predictive neurons were active both on hit and false alarm trials, so that their activity was better modulated by the animal’s motor response than by the identity of the GO or NOGO sequence (Figure 3E). Moreover, their responses were temporally locked to licking rather than to stimulus presentation (Figure 3E). In these neurons, responses preceding licks were not stereotypical but differed for early and late licks in a trial, reflecting the licks’ relevance during task engagement (Figure 3E, right inset). This differentiated such action predictive neurons from a rarer set that appeared purely to reflect the motor action of a lick, regardless of context (termed ‘licking’ in Table 1).

Finally, some neurons displayed further readily identifiable preferences for trial-to-trial task outcome variables, including reward delivery (Figure 3F) and reward prediction error (Figure 3G).

Overall, across our data set of S1bf neurons in well-trained mice (n = 315 neurons), n = 179 (57%) were classifiable according to the categories above (Table 1). Within the classifiable neurons, those with a pure sensory response were in the minority (n = 54; 30%). Moreover, only a single neuron was identifiable as providing pure sensory categorical encoding of trial type (i.e. responding to either the GO or NOGO sequence in a manner independent of the animal’s behaviour, as in the ‘sensory category’ hypothetical example in Figure 3C). In contrast, we frequently observed action predictive neurons whose responses reflected the learnt association between a stimulus perceived as the target sequence, and the consequent action (n = 75; 42%).

### Diverse neuronal response types in well-trained S1bf

The visual classification described above suggested that S1bf neurons in well-trained animals behaved heterogeneously in the context of the task. To assess this quantitatively, we analyzed the extent to which different cells preferentially responded with distinct patterns or profiles (as reflected in Figure 3C-G). If neuronal response profiles during a trial fell into distinct subsets, this would be reflected into a degree of response similarity between subsets of neurons greater than that expected if their profiles varied at random. To test for this, we performed a projection angle index of response similarity (PAIRS) analysis (Methods) [40]. We found that the median response similarity between neurons in our data set, given by this PAIRS measure, was greater than for any of 10000 random surrogate neuronal data sets (Figure 4A; p < 10^−4^). Thus, PAIRS analysis established that neuronal responses clustered more than expected by chance.

**Figure 4.**
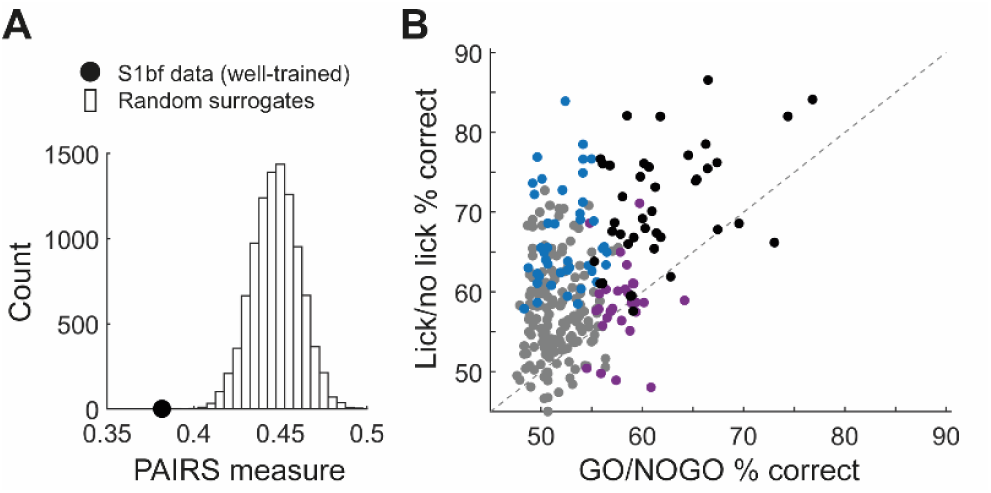
Diverse response types in well-trained S1bf. **(A)** Median PAIRS value (index of response similarity) for experimental S1bf data set compared to distribution for 10000 random surrogates. The response properties of experimental neurons clustered more than expected by chance. **(B)** Classification performance (% correct) supported by individual neurons. Data points shows lick vs no lick performance plotted against GO vs NOGO performance for each neuron. Black dots: neurons with significant performance on both lick vs no lick and GO vs NOGO. Blue: neurons with significant performance on lick vs no lick. Purple: neurons with significant performance on GO vs NOGO. Grey: neurons with significant performance on neither. Dashed line: equality.

This result of PAIRS analysis, taken together with the observation that neuronal responses reflected distinguishable aspects of the task (Figure 3D-G), suggested the existence of neurons with distinct functional properties. We thus developed a classifier analysis to quantify whether a neuron conveyed information about sensory trial type (GO vs NOGO) or the animal’s response (lick vs no lick), based on how the neuron’s response evolved during a trial (Methods). In brief, we used the disparity between the ΔF/F_0_ time course over trials of different types to decode whether the trial was either GO vs NOGO or lick vs no lick.

This analysis showed that 25% of neurons (70 out of n = 277) were able to support classification of sensory trial type (GO vs NOGO) to criterion level (Figure 4B; defined as the neuron performing better than 95% of surrogate classifiers constructed by shuffling trial labels). Note that this proportion does not include all neurons responsive to sensory stimulation: for example, a ‘sensory cue’ neuron such as the one in Figure 3D would not be able to classify GO vs NOGO as it would respond very similarly on both types of trial. In terms of classifying whether the mouse response on a trial was lick vs no lick, 30% of neurons (86 out of n = 284) could perform to criterion level (Figure 4B).

Figure 4B plots classification performance on lick vs no lick against performance on GO vs NOGO, shown for all neurons for which it was possible to compute both classifiers (n = 272). Across neurons, classification performance was higher for lick vs no lick than for GO vs NOGO (p < 10^−34^, Wilcoxon signed rank test). Notably, multiple neurons supported lick vs no lick classification with a high level of performance (∼70% or above), consistent with our identification of action predictive neurons in the observations above. While some neurons could discriminate the GO vs NOGO sequence with good performance, rarely did they do so without also showing sensitivity to the upcoming lick action. In other words, sensory representations in individual S1bf neurons of well-trained animals were tangled with, and not independent from, action representations. This is a striking result considering that S1bf is a textbook sensory area, and deviates from the expectation that sensory responses would dominate our data set.

### Neuronal responses in S1bf selective to the target sequence and associated actions appear as soon as the association is learnt

Given that a salient feature of our well-trained S1bf data was the presence of neurons whose activity predicted licking to the learnt sequence or was sensitive to trial outcome (Figure 3), we sought to determine when these responses arose during task learning. We therefore repeated the above analyses in animals that had already been trained to remain head-fixed but not yet to detect or recognize a sequence stimulus (n = 4 mice; Figure 5). Imaging took place starting from the first training session and up to the fourth. Mice began to lick selectively to the GO as compared to the NOGO stimulus in the second session.

**Figure 5.**
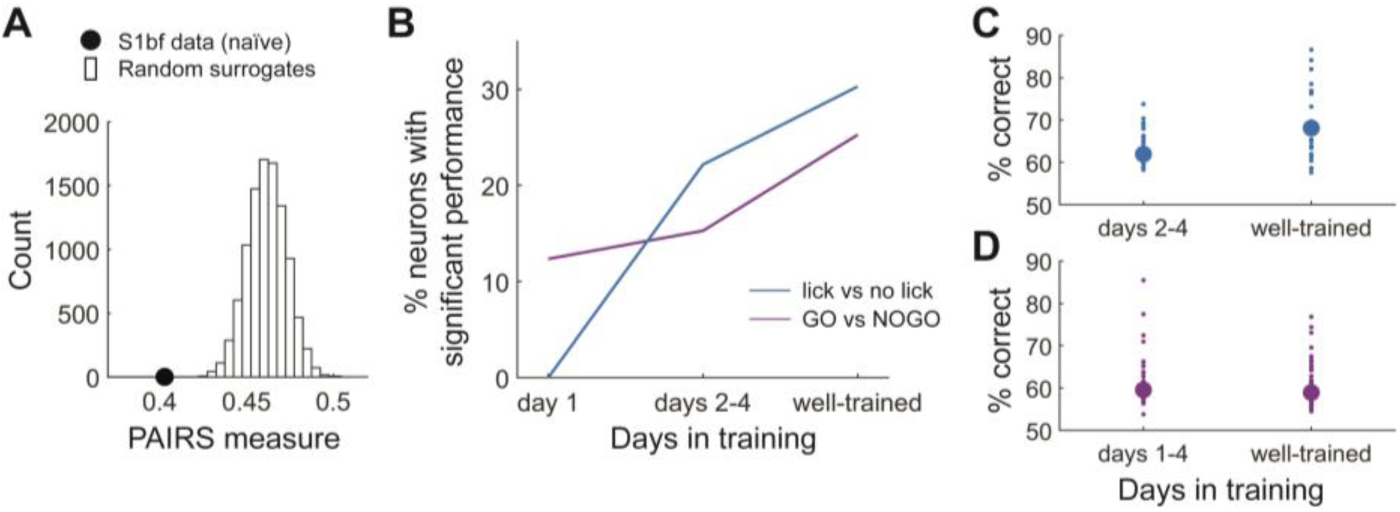
Progression of neuronal selectivity to sequence identity and the associated actions over the course of training. **(A)** Median PAIRS value (index of response similarity) for naïve S1bf data compared to distribution for 10000 random surrogates. **(B)** Progression of the percentage of neurons with significant performance with days in training. Progression is significant for both curves: for GO vs NOGO classification, χ ^2^ statistic = 11.36, p = 0.0034, χ^2^ test. For lick vs no lick, χ ^2^ statistic = 9.97, p = 0.0068, χ ^2^ test. **(C)** Classification performance for significant neurons on lick vs no lick, for different stages of training. Small dots: individual neurons. Thick dot: median. **(D)** Classification performance for significant neurons on GO vs NOGO, for different stages of training. Small dots: individual neurons. Thick dot: median.

We found that neurons in these naïve animals participated more sparsely in task encoding: a smaller proportion than in well-trained mice could be readily classified into the categories defined earlier (Table 2; 20% of 356 neurons vs 57% of 315 neurons; p < 10^−22^, odds ratio 5.19, Fisher’s exact test). For example, we found no action predictive neurons at all in the first training session. Thus, training resulted in a denser representation of sensory and task-related variables at the population level.

**Table 2.** Proportions of visually identified neurons responding to sensory and task variables in naïve S1bf.

| Sensory cue | Sensory feature selective | Sensory category | Decision/action predictive (during stimulus period) | Decision/action predictive (during response period) | Licking | Reward delivery and error | Unclassified |
| --- | --- | --- | --- | --- | --- | --- | --- |
| 23<br>(6.5%) | 22<br>(6.2%) | 2<br>(0.56%) | 7<br>(2%) | 4<br>(1.1%) | 4<br>(1.1%) | 10<br>(2.8%) | 284<br>(80%) |

PAIRS analysis showed that, as in well-trained animals, S1bf neurons in naïve mice also had clustered response profiles (Figure 5A; p < 10^−4^). Accordingly, we again performed a classifier analysis to quantify the fraction of neurons that could support classification of trial identity (GO vs NOGO) or licking response (lick vs no lick).

The ability of neurons to classify trial type progressed over the course of training (Figure 5B). For neurons imaged in the first training session, 12% (11 out of n = 89) could support classification of GO vs NOGO trial identity. In naïve animals overall, 14% of neurons (46 out of n = 318) could classify GO vs NOGO trial type; this fraction was significantly lower than in well-trained animals (25%; p = 0.0081, odds ratio 1.75, Fisher’s exact test), consistent with the notion that the proportion of neurons representing trial type increased as a result of training (i.e. the population sparseness decreased), as described above.

Classification performance on lick vs no lick trials progressed in a slightly different manner: in the first training session we found no neurons that could support the lick vs no lick classification (Figure 5B). This changed from the second session: 22% of neurons imaged in sessions 2-4 (39 out of n = 176) could classify trials as lick vs no lick, a percentage not significantly different from that in well-trained animals (30%; p = 0.174, odds ratio 0.732, Fisher’s exact test). Thus, neurons whose activity differed significantly on lick vs no lick trials were first found on the second day of training, at the same time as animals first associated the target sequence with licking for a reward and the non-target sequence with suppression of licking.

The trial-by-trial classification performance of significant ‘lick vs no lick’ neurons increased during training (Figure 5C; n = 39 for sessions 2 to 4, n = 86 for well-trained, p = 0.00019, Wilcoxon rank sum test). In contrast, although the proportion of significant ‘GO vs NOGO’ neurons increased with training (Figure 5B), their classification performance did not (Figure 5D; n = 46 for sessions 1 to 4, n = 70 for well-trained, p = 0.168, Wilcoxon rank sum test). Thus, neurons whose activity reflected sensory trial type did not refine this selectivity with training.

These results demonstrate that, as a result of the explicit learning of a target sequence of whisker stimulation, neurons in somatosensory cortex reflect multiple task variables, and particularly the learnt association between the sensory sequence and the goal-directed decision to act.

## Discussion

Training mice on a whisker-mediated elementary sequence discrimination task, we found three main results: (i) animals appear to solve the task by seeking the earliest cues that predict the identity of the target sequence, but perform better when integrating or deliberating over this information; (ii) somatosensory cortex (S1bf and S2) is needed for performing the task; and (iii) neurons in superficial layers of S1bf display heterogeneous selectivity to task variables including sensory input, the animal’s action decision, and trial outcome – with the latter selectivity being acquired as a result of learning to associate the target sequence with the action needed for reward. In naïve animals, presenting a target sequence activates neurons selective to sensory input; upon training, presenting the same (rewarded) target sequence activates neurons that now, in large numbers, predict the animal’s learnt licking response. These results defied our expectation that neuronal activity in S1bf of well-trained animals would primarily represent features of sensory input, perhaps refined by learning; rather, we found neurons that embody learnt associations between sequences and the corresponding behaviour, and predict expected actions and outcomes in the context of the task.

### S1bf and S2, but not PPC, participate in goal-directed tactile sequence recognition

Our optogenetic experiments showed that suppressing activity in S1bf and S2 interferes with whisker-mediated sequence recognition, in a manner consistent with an interruption in the flow of sensory information. This indicates that S1bf and S2 are needed to perform the task. Similar conclusions have been reached for other tasks that demand recognition of whisker input streams [34, 41-45]. In contrast, S1bf has been shown to be unnecessary for simple tasks involving detection of whisker motion [18, 19], in a manner reminiscent of the residual function observed in multiple senses after cortical lesions or removal [46-49]. While sensory cortex is not required for stimulus detection or localization, it may be needed for organizing sensory data into objects or concepts according to previous experience, including context-dependent change detection [50, 51]. The effect of S1bf and S2 suppression was specific to these somatosensory centres. Interfering with PPC activity had no discernible effect on task performance; we note that under our task design, no comparisons of temporally separate stimuli were required and the stimulus sequence did not need to be kept in working memory.

### Plasticity elicited by goal-directed learning of a sequence couples sensory neurons to action decisions

Neurons in the naïve S1bf have limited temporal integration: they report on, or encode, sensory input accumulated over just a few tens of milliseconds [21-26]. In earlier studies, this temporal integration remained limited even after learning of tasks in which performance could benefit from greater integration [25]. This property of being responsive to sensory signals that are current rather than accumulated over time is shared with neurons in the primate primary somatosensory cortex [52], suggesting a common principle across tactile pathways.

Neurons can intrinsically discriminate between stimulus sequences, and modelling studies have shown that generally observed forms of synaptic plasticity can endow neurons with sensitivity to input patterns that last longer than the neuron’s membrane integration timescale (e.g. [53-55]). Our expectation was therefore that conditioning on a specific sequence might produce S1bf neurons that preferentially represented that sequence, now habitually present in the animal’s life and associated with a desirable goal. This would fit in with a framework in which the role of sensory cortex is primarily to provide sensory representations [56] and learning refines these representations to become more predictive of upcoming sensory signals or increasingly modulated by behaviour [57]. Instead, we found neurons that – after training but not at the outset of training – directly embodied the association between the presence of the target sequence and the appropriate learnt response, but did not respond selectively to the sequence independently of the animal’s response. We suggest that these neurons may have become predictive of the learnt outcomes associated with sensory stimuli to which they were originally responsive [51]. Note that upon learning there remained a considerable number of other neurons that responded simply to the onset of the sensory stimulus (Figure 3C, D), and these showed no modulation related to licking; thus, our results are inconsistent with sensory signals being occluded by a spread of prevalent lick-related activity.

### Non-sensory representations in sensory cortex

The classic account of perception posits a serial feedforward scheme whereby successive processing steps map neatly onto neuronal activity elicited in distinct brain regions, with each region’s neuronal responses being classified as essentially sensory, decision-or action-related. These conclusions were originally derived from classic experiments carried out under anaesthesia; in this condition, neurons in the primary sensory cortex respond to specific physical features of stimuli, while neurons in higher areas such as the prefrontal cortex (PFC) cannot be characterized as sensory feature selective. More recent experiments have often involved training animals to produce a sensory-guided response only after the period of sensory stimulation has concluded. This temporal separation between stimuli and action readily allows for isolation of neuronal activity reflecting different aspects and stages of the trained behaviour, and has produced deep insights into how these map onto neuronal processing stages (reviewed in e.g. [52]).

Conversely, our results show that in situations where an animal is permitted to act while an ongoing stimulus is still being presented, a complex interplay occurs between brain regions, and neuronal activity propagates in an intricate pattern, reflecting many aspects of the task. Neurons in the primary sensory cortex respond to multiple task variables, including not just sensory signals but also the animal’s decision on whether to act and the decision’s outcome (e.g. the presence or absence of an expected reward). This rich pattern of activity is likely to represent the neuronal substrate for the animal’s interactions with a dynamic environment as often played out in real life, where sensory signals and our responses to them are ongoing and intertwined in time, rather than separate.

Although inconsistent with a classic feedforward picture of cortical processing, our data are consistent with findings gathered by multiple laboratories in other contexts. In those studies, variables affecting neurons in sensory cortex include the decision to act and choice of action; arousal and attention; spontaneous gestures and motions, and consequent mismatches between actual and expected sensory input; motor activation relevant to task execution; and expected and actual rewards and their timing [34, 35, 58-85]. In our data, non-sensory parameters do not just modulate the responses of sensory cortex neurons to a target stimulus; rather, in a subset of neurons, the responses reflect action decisions or reward outcomes instead of sensory cues or features, consistent with other recent studies [79, 86, 87].

Sensitivity to multiple sensory and behavioural variables is a well-established property in higher cortical areas such as PFC [88-92] and parietal cortex [40, 93]. The findings above suggest that, upon learning, a similarly rich representation becomes shared by early sensory areas. This could happen through top-down connections [94, 95] broadcasting a “copy” or version of frontal cortex responses to sensory cortex. Supporting this idea, non-sensory activity reflecting motor actions spreads widely across the dorsal cortex when an animal is engaged in a learnt task [83, 96-100], perhaps as an “efferent copy” signal reflecting the animal’s behaviour [101]; this broadcast seems to originate in a frontal premotor region linked to action selection and motor preparation [45, 96-98, 100, 102]. Activity related to decisions is prominent in PFC, premotor cortex and the basal ganglia (dorsal striatum): these areas have been linked to evidence accumulation in favour of one or another sensory stimulus, to categorisation, and to reaching the decision itself [28, 52, 100, 102-105]. In rodents, all of these areas receive direct connections from the somatosensory cortex, and learning an association between sensory information and specific actions is likely to reinforce feedback loop interactions between sensory and higher regions [33, 34, 74, 86, 98, 106]. The links between a stimulus and the appropriate action could be learned initially in one of the higher areas; depending on familiarity with the task and behavioural context, top-down connections to S1bf and S2 could then help link the lower-level representation of the target stimulus to its consequent action. It will be important to understand the circuit plasticity mechanisms underlying this process.

How to interpret the action-predictive neurons we found in sensory cortex? These were present in trained but not naïve animals. We surmise that they may have originally been selective to stimulus features present in the target sensory sequence, and then, by virtue of being active at an appropriate time during target presentation, ‘tagged’ during learning as being able to participate in driving the goal-directed response prompted by the target sequence. By a process of associative synaptic potentiation, these neurons might then have become more strongly connected to postsynaptic partners capable of affecting behaviour. Results from the auditory modality suggest plasticity phenomena consistent with this account: neurons in the auditory cortex sensitive to a frequency range present in a target stimulus eventually become able to drive the GO response [51, 73, 104, 107]. In another whisker-mediated task, in which mice learn to lick in response to a detected whisker deflection, S2-targeting neurons in S1bf also acquire responses predictive of licking [74].

An alternative interpretation is that the responses of action-predicting neurons in sensory cortex simply reflect the top-down broadcast of an action choice from higher decision-making centres. If this interpretation is true, it raises the question of how the sensory information needed to reach the decision is relayed to higher areas: our data suggest that only a small minority of neurons in trained S1bf convey purely sensory information. Unravelling this issue will be important for understanding the specific pathways converting sensory input into a sensory-guided, goal-directed decision [52].

### Experimental considerations

Our findings underscore the importance of measuring behavioural output as well as sensory input to interpret neuronal responses, and of including error trials in the analysis. Without monitoring the timing of individual licks it would be impossible to distinguish a categorical sensory neuron from an action predictive neuron. Conversely, the present GO/NOGO experimental design does not allow dissociation between the decision to act and the specific choice of action, and thus does not allow us to distinguish between action predictive and choice predictive activity [100, 108].

We based our analyses of two-photon recordings on the fluorescence (ΔF/F_0_) time series rather than on activity reconstructed by deconvolution. The results of deconvolution methods can be highly sensitive to parameter choice, giving rise to variable conclusions as to neuronal response properties [109, 110]. On the other hand, any analysis based on ΔF/F_0_ will overestimate neuronal correlations and give estimates on neuronal encoding of task variables that are likely to be a smoothed, filtered version of that occurring in reality [110, 111]. Accordingly, our conclusions on clustering of neuronal response properties are not built on an absolute estimate of correlations between responses, but on assessments of correlations relative to a random surrogate version of activity. Any distortions of response temporal patterning resulting from calcium imaging would tend to smear out differences between neurons, so our assessment of neuronal selectivity is likely to underestimate the true amount of tuning heterogeneity.

In our findings, learning superimposed behavioural associations onto the responses of neuronal populations in a sensory area. Cortical populations, even those located at early stages of processing, modulate their responses depending on behavioural context, and neuronal codes for task parameters depend on the specifics of the task. It is likely that the specific representations of task parameters uncovered here reflect the nature of the animals’ training: specifically, the fact that mice learned to explicitly recognize a particular sensory sequence. When an animal is exposed to repeated sequential sensory patterns but not conditioned on them, i.e. is not asked to learn an explicit relationship between the patterns and a goal-directed motor action, changes occurring in primary sensory cortex may be different and include prominent refinements in the sensory tuning of neurons, leading them to enhance categorization [112] and potentially become sensitive to sequence structure over extended periods of time [113-116]. Understanding how neurons distinguish between behaviourally relevant, but not explicitly taught, sequences will be important to illuminating our ability to recognise, make sense of, and value ongoing patterns in our surroundings.

## Methods

All procedures were conducted in accordance with national (UK Animals (Scientific Procedures) Act 1986) and international (European Union directive 2010/63/EU) regulations for the care and use of animals in research. Personal and project licenses to carry out the work were approved upon institutional and Home Office review. Experimental mice were males on a C57BL/6J background, 4-9 weeks old at the time of surgery and bred at the University of Sussex.

### Surgical procedures

Details of head bar implantation surgery have been published elsewhere [16, 117]. Briefly, under aseptic conditions, mice were anaesthetised using 1.5-2.5% isoflurane in O_2_ and placed into a stereotaxic apparatus (Narishige, Japan) with ear bars previously coated with EMLA cream. We monitored anaesthetic depth by checking spinal reflexes and breathing rates. Body temperature was maintained at 37°C using a homeothermic heating pad (FHC). Eyes were treated with ophthalmic gel (Viscotears Liquid Gel, Novartis, Switzerland) and the entire scalp washed with povidone-iodine solution. An area of skin was removed (an oval of approximately 15 mm x 10 mm in the sagittal plane) such that all skull landmarks were visible and sufficient skull was accessible to securely fix a titanium or stainless steel head bar. The exposed periosteum was removed and the bone washed using saline solution, dried with sterile swabs and then scraped with a scalpel blade to aid bonding of glue. Cyanoacrylate glue (Vetbond, 3M) was applied to bind skin edges to the skull and as a thin layer across the exposed skull to aid bonding to the dental acrylic. A custom titanium or stainless steel head bar (dimensions 22.3 × 3.2 × 1.3 mm; design by Karel Svoboda lab, Janelia Farm Research Campus, Howard Hughes Medical Institute, https://hhmi2.flintbox.com/public/project/26526/) [117] was placed directly onto the wet glue centred just posterior to lambda. Once dry, we scraped the glue surface to improve bonding and fixed the head bar firmly in place by applying dental acrylic (Ortho-Jet, Lang Dental) to the head bar (on top and behind) and the skull (anterior). Mice were given buprenorphine (0.1 mg/kg, I.P.) and further EMLA cream to the paws and ears. Once the acrylic was set, anaesthesia was turned off and animals returned to the cage. On the day of surgery and for the next two consecutive days 200 μl of non-steroidal anti-inflammatory drug (Metacam oral suspension 0.5mg/ml; Boehringer Ingelheim) was mixed with food pellets soaked in water until they became mash. Animals were housed individually and allowed to recover for one week post-surgery, with health and weight monitored daily.

### Housing and training

Animals were housed in cages with bedding, tubes, running wheels and a plastic plate attached to a custom platform used to provide daily water, and kept on a reverse 50:50 light-dark (LD) cycle.

#### Water control

To motivate mice to perform the task we employed a water restriction protocol [117] and made water available as a reward for correct recognition of GO stimuli. Mice cope well physiologically with water restriction, as they are adapted to life in semiarid environments [118]. Dry food was available at all times. We observed a mild increase in motivation when mice were given sunflower seeds before tasks.

Mouse water intake was regulated so that animals were motivated to perform for 200 or more trials per session under our conditions (45-55% humidity, 20°C and atmospheric pressure; reverse 50:50 LD cycle), while remaining active and healthy. The water control protocol started 7 days after head bar implantation surgery. A single training session was performed on each day when training was carried out. Animals received water during the training session; reward water intake was determined by weighing the animal before and after the session together with collected faeces, and was typically 0.1-0.4 ml. Mice were then given further ad libitum water during a finite (usually 1 min) free drinking period after the end of the session. On days with no training, mice were given free access to 1.5 ml, which corresponds to 50% of average ad libitum water intake for C57BL/6J mice (Mouse Phenome Database from the Jackson Laboratory: http://www.jax.org/phenome). The health of animals under water restriction was assessed daily (dehydration, weight, grooming, movement) and a checklist filled. Mice initially lost weight but then increased body mass gradually over the course of water restriction. Sensory discrimination training began after 9 days on water control.

#### Animal handling and training set-up

Mice were trained to enter a head fixation device using a shaping procedure. We initiated water control one week after head bar implantation. On days 1 and 2 animals were given 1.5 ml of water placed in their cage. On days 3 and 4 animals were introduced to the experimenter. They were first left to smell and explore the experimenter’s hand while in their cage, then gently picked up using a tube and returned to the cage several times while given sunflower seeds and water from a syringe. On days 5 and 6 mice were introduced to the head fixation device. They received sunflower seeds and water via a syringe only when inside the device (but not head-fixed). At this stage, mice were grooming and eating in the head fixation apparatus without any signs of distress. On days 7 and 8 animals were given a sunflower seed and after ingestion were head-fixed and given water via a syringe. Animals became accustomed to head fixation and expected to receive water from the spout situated in front of their head. On day 9, mice began the task. Animals were trained in the dark; illumination, if necessary, was provided by a red lamp.

We used two device designs. One design consisted of an acrylic tube (32 mm internal diameter) with its head end cut to enable access to the implanted head bars. The tube was placed on Parafilm or a rubber glove and clamped into a v-shape groove. This support acted to stabilise the tube, collect faeces and prevent mice from grasping stimulus apparatus and the lickport. The second design consisted of a custom 3D-printed treadmill on which mice could locomote freely (design by Leopoldo Petreanu, Champalimaud Centre for the Unknown). A metallic mesh was fixed over the treadmill to surround the mouse’s body, allowing the animal to feel comfortably enclosed rather than exposed. The ends of the head bars were inserted into notches on two head fixation clamps and tightened using thumbscrews.

Water rewards were provided through an electrical lickport ending in a spout made from a blunted gauge 13 syringe needle. Water flow from an elevated container was controlled via a solenoid valve (LDHA1233215H, The Lee Company, France). The acrylic tube was lined with aluminium foil. Terminals from an A/D input of a signal processor were connected to the water spout and the foil or the metallic head bar holder, so that tongue contacts with the lick port created brief elevations in voltage consistent with lick durations. This opened the solenoid valve for an adjustable amount of time, delivering 1-2 μl of water. Correct positioning of the lickport was an important aspect of training: in the first sessions it was placed relatively close to the mouth of the animal, to facilitate initial successful collection of rewards, but was gradually moved away from the mouth during training to avoid development of impulsive licking.

#### Stimulus design and delivery

Stimulus sequences were constructed in Matlab (Mathworks, USA). Stimulus playback and trial control was performed either via a signal processor (RP2.1, TDT, USA) controlled with code written in ActiveX, or via a Bpod/PulsePal (Sanworks LLC) open-source Arduino-based system [119] controlled with code written in Matlab. Trial outcomes were recorded in Matlab.

Multiple whiskers on one side of the snout were trimmed to 1 cm and placed into a 10 mm^2^ metallic mesh grid (at least 3 whiskers in the grid), glued to a piezoelectric actuator (PL127.11, Physik Instrumente, Germany) and positioned ∼1 mm from the animal’s fur.

To ensure that mice detected the lowest amplitude filtered noise part of the sequence stimuli (Figure 1B), animals (n = 3) that had successfully learned normal detection were trained to detect the lowest amplitude syllable. All of the animals accomplished high performance on detection within a single session (mean 81% correct, SD 7.83%; n = 4 sessions).

### Optogenetics

To suppress activity in dorsal cortical areas, we photostimulated channelrhodopsin-2 (ChR2) in GABAergic interneurons of VGAT-ChR2-EYFP mice (Zhao et al., 2011) (breeding pairs from The Jackson Laboratory; stock number: 014548). Three mice were used to measure light transmission through the clear-skull cap preparation, one mouse to verify expression of ChR2, and six mice to characterize photoinhibition. Of these, four were used for behavioural training and optogenetics experiments.

#### Surgeries

Mice were implanted with a head bar as described above. At the same time, a clear-skull cap was added [45, 100, 120], as follows. After marking bregma using a surgical marker, covering the bone surface with a thin layer of cyanoacrylate glue and allowing the glue to dry, two to three thin layers of UV curing optical adhesive (Norland Optical Adhesives #81, Norland Products Inc.) were applied to the skull and cured using a UV LED (DC4100, Thorlabs). Next, the headbar was attached to the skull and fixed with dental cement. To avoid scratches during the animal’s recovery and training period and keep the clear-skull cap’s surface smooth, it was covered with a silicon sealant (Kwik-Cast, World Precision Instruments). The sealant was removed prior to each experiment and a new layer applied before the animal returned to its cage.

#### Photoinhibition

Light from a 473 nm modulated diode laser system (Cobolt 06-MLD, Laserlines) was controlled with digital modulation (< 2.5 ns rise time). The laser head was fibre coupled (FC/PC) to a 2 m length multi-mode optical fibre (200 μm diameter, Laserlines). Light coming out of the fibre was collimated using an adjustable collimator (350-700 nm, CFC-8X-A, Thorlabs) and passed through a coated plano convex lens (LA1951-A, Thorlabs), to be focused onto the surface of the clear-skull cap. Light modulation followed a 50 Hz square wave control signal generated by a voltage pulse generator (PulsePal, Sanworks). Laser power was calibrated using a handheld power meter (NT54-018, Edmund Optics). The laser beam had a Gaussian profile (FWHM 364 μm), determined using a CMOS camera (DCC1545, Thorlabs; pixel size 5.2 μm) and analysed with Fiji.

Light transmission through the clear-skull cap was measured on a separate group of mice that underwent the preparation surgery and were then euthanized (n = 3 mice). The clear-skull cap (skull, cyanoacrylate glue and UV curing optical adhesive) was next isolated and laser power measured before and after passing through it. Light transmission was 36 ± 2% (SD). After calibration, laser power was set to approximately 3.4 mW at the brain surface.

At the beginning of each experiment, the mouse was head-fixed and silicon sealant removed. The laser beam was positioned over bregma and subsequently moved to the brain area of interest with a motorized manipulator (MP-225, Sutter Instruments). A single area was perturbed in each session.

The distances between optogenetic stimulation sites in S1bf, S2, PPC and wM1 were at least 1.2 mm, > 3x the FWHM of the laser beam. Note that our stimulation sites in wM1 and S1bf, which yielded contrasting effects (Figure 2B, E), were closer together than those in S1bf and S2, which yielded similar behaviour (Figure 2B, C). Thus the similarity between S1bf and S2 effects was not explained by site overlap.

### Two-photon imaging

#### Surgeries

Thy1-GCaMP6 mice expressing GCaMP6f in pyramidal neurons [36, 121] (founder lines GP5.5 and 5.17) were implanted with a head bar as described above. A circular 3 mm diameter craniotomy was made to expose the brain. A cranial window, consisting of a 3 mm circular coverslip and a 5 mm circular coverslip (Harvard Instruments), was placed over the craniotomy and secured in place with cyanoacrylate tissue sealant (Vetbond, 3M). Following recovery, mice were trained to perform the task while head-fixed under the two-photon microscope, using a shaping procedure as described above. On concluding the experiments, we checked for specificity of GCaMP6f expression in excitatory neurons by staining Thy1-GCaMP6 mice for VGAT expression (mean 1.4% of Thy1-positive neurons expressed VGAT, range 1.0-1.9%; n = 4 mice).

#### Imaging

A two-photon microscope with galvanometric scanning (Scientifica) was controlled by Scanimage software (Vidrio Technologies). Illumination was provided by a Ti:sapphire Chameleon Vision S laser (Coherent Technologies) tuned to 940 nm and focused through a 20x/1.0NA water immersion objective (Olympus). Laser power under the objective was 100-120 mW. Frame scanning (256×100 pixels) was performed at 10.8 Hz.

#### Image processing

Image processing was carried out using Suite2p [122] running in Matlab. After registration and motion correction, ROIs were automatically detected and manually adjusted. Raw fluorescence was extracted for each ROI and corrected for neuropil contamination (F = F_raw_ – αF_neuropil_) [121, 123]. Baseline fluorescence F_0_ was computed using a 2-3 min sliding window, using the 5^th^ percentile of the raw distribution within the window for highly skewed cells, or the median for cells with a symmetric distribution [37].

### Behavioural analysis

We quantified behavioural performance as in [16], using a percent correct metric determined over a 50-trial sliding window during the course of a session and corrected for the proportions of GO and NOGO trials. To obtain an upper bound on average decision times in a session, we determined when, on average, the lick rates for GO and NOGO trials began to diverge during the course of a trial (discriminative lick latency, DLL) [16]. We first subtracted the lick rate curve for NOGO trials from that for GO trials, and set a threshold for when this subtracted curve became positive (i.e. when GO licks surpassed NOGO licks) by determining the 95% confidence limit for the distribution of subtracted lick rate curves throughout the trial for 50 randomly subsampled sets of GO and NOGO trials (time points sampled at 100 ms resolution).

### Neuronal response analysis

For each neuronal data set (S1bf in well-trained and naïve animals) we computed PAIRS analyses separately. To do this, we represented each neuron in the data set by a vector of length 246, constructed by concatenating the cell’s average ΔF/F_0_ responses to the four trial types with reference to trial time together with the hit and false alarm trials referred to first lick time. We computed 8 principal components (PCs), which captured over 90% of the variance in the data, and recast each neuron in terms of a response feature vector consisting of the projections of its average response vector onto the 8 PCs. To evaluate response similarity between any two neurons, we took the dot product between their 8-dimensional response feature vectors, which was the cosine of the similarity angle *θ* between them. For each neuron z, we then obtained the mean of this angle with the neuron’s k nearest neighbours, *θ*_*z*_^*(k)*^. The PAIRS measure was computed as the median of *θ*_*z*_^*(k)*^ across all neurons in the data set, represented as a thick dot in Figures 4A and 5A. To determine whether the response similarity given by the PAIRS measure differs from that expected if occurring by chance, we generated 10000 surrogate neuronal data sets. For every surrogate neuron, the value of each component of the 8-dimensional feature vector was drawn at random from its empirical distribution across real neurons. The plots show results computed for a choice k = 3 of the number of nearest neighbours [40]; results were consistent for a wide range of choices of k (3-8).

To check whether the significant clustering present in PAIRS data might be influenced by neurons being grouped according to their session or mouse, we carried out PAIRS analyses separately on data collected from individual experiments, including only sessions where over 25 neurons were imaged. A common PCA basis was used for all experiments within the same category (well-trained or naïve). For all the individual experiments, the experimental PAIRS measure was smaller than the surrogate ones at greater than 95% confidence level.

To estimate how well the activity of a neuron could support classification of trial type (GO vs NOGO, or lick vs no lick), we trained a separate support vector machine (SVM) for each neuron, using as input the ΔF/F_0_ time course from all trials, with each labelled according to the trial type of interest. To limit bias, if the proportion of trials with licks was more than 75% (a common occurrence in early training), we randomly removed lick trials until the proportion was better balanced than 75%/25%; we only used neurons for which > 100 trials remained after this operation. Training was performed using the sklearn SVM library and classification performance assessed using 5-fold cross validation.. A linear kernel was used, and several regularisation parameters tried (.001, 0.01, 0.1, 1) before choosing the one with best cross validated score. The training procedure was repeated for 100 surrogates generated by shuffling trial labels. Neurons were deemed to support classification at a significant level of performance if they performed better than 95 of the surrogates. We repeated this analysis using information theory methods by evaluating the information about trial type conveyed by each neuron’s trial-by-trial response, and obtained results qualitatively identical to those based on classifiers.

## Acknowledgments

This work was supported by the UK Medical Research Council (grant number MR/P006639/1) and the University of Sussex internal research development fund. The authors declare no competing financial interests. We thank V. Jayaraman, R. Kerr, D. Kim, L. Looger, K. Svoboda and the HHMI Janelia Farm GENIE project for mice expressing the genetically encoded calcium indicator GCaMP6f; Simon Peron for help with the head bar adaptation and cranial window construction; Matteo Carandini and Charu Reddy for advice on the clear skull-cap preparation; Chen Qian for help with the optogenetics set-up; Moira Eley for help with animal maintenance and training; and members of the Maravall lab for comments on the manuscript.

